# Combining high-throughput micro-CT-RGB phenotyping and genome-wide association study to dissect the genetic architecture of tiller growth in rice

**DOI:** 10.1101/247841

**Authors:** Di Wu, Zilong Guo, Junli Ye, Jianxiao Liu, Guoxing Chen, Jingshan Zheng, Dongmei Yan, Xiaoquan Yang, Xiong Xiong, Qian Liu, Zhiyou Niu, Lizhong Xiong, Wanneng Yang

## Abstract

Traditional phenotyping of rice tillers is time consuming and labor intensive and lags behind the rapid development of rice functional genomics. Thus, dynamic phenotyping of rice tiller traits at a high spatial resolution and high-throughput for large-scale rice accessions is urgently needed. In this study, we developed a high-throughput micro-CT-RGB (HCR) imaging system to non-destructively extract 730 traits from 234 rice accessions at 9 time points. We used these traits to predict the grain yield in the early growth stage, and 30% of the grain yield variance was explained by 2 tiller traits in the early growth stage. A total of 402 significantly associated loci were identified by GWAS, and dynamic and static genetic components were found across the nine time points. A major locus associated with tiller angle was detected at nine time points, which contained a major gene *TAC1*. Significant variants associated with tiller angle were enriched in the 3'-UTR of *TAC1.* Three haplotypes for the gene were found and tiller angles of rice accessions containing haplotype H3 were much smaller. Further, we found two loci contained associations with both vigor-related HCR traits and yield. The superior alleles would be beneficial for breeding of high yield and dense planting.

**Highlight:** Combining high-throughput micro-CT-RGB phenotyping facility and genome-wide association study to dissect the genetic architecture of rice tiller development by using the *indica* subpopulation.

## Introduction

Rice is one of the most important food crops both in China and worldwide (Zhang, 2008). Selecting plants with the ideal tiller structure is a key issue for domesticating rice and improving its yield (Wang *et al*., 2008). With the rapid development of functional genomics and molecular breeding, rice researchers and breeders often need to screen thousands of lines in a short time for the targeted phenotypic traits under different growth conditions (Fiorani and Schurr, 2013). However, traditional phenotyping, particularly tiller measuring, is time consuming and labor intensive and lags behind the development of rice genomics (Houle *et al*., 2010; Furbank *et al*., 2011). To bridge the gap, progress in high-throughput phenotyping technology is required to accelerate gene discovery and rice breeding (Huang *et al*., 2013; Spalding *et al*., 2013).

Over the past 20 years, many non-destructive and high-throughput phenotyping methods have been constructed to obtain plant phenotypic data. These methods include shoot phenotyping platform in greenhouse such as TraitMill (Reuzeau *et al*., 2005), PHENOPSIS (Bacmolenaar *et al*., 2015), Phenoscope (Sébastien *et al*., 2013), Scanalyzer 3D (Junker *et al*., 2014), root phenotyping in greenhouse such as GROWSCREEN-Rhizo(Nagel *et al*., 2012), GiA Roots and Rootowork (Topp *et al*., 2013), field phenotyping platform such as BreedVision (Busemeyer *et al*., 2013), and unmanned aerial vehicles (Berni *et al*., 2009). With rapid progress in photonics, several novel imaging techniques have been adopted in crop phenotyping. These techniques include near-infrared imaging to estimate plant disease (Bock *et al*., 2010), stereo camera systems to quantify rape leaf traits (Xiong *et al*., 2017), fluorescent imaging to diagnose biotic or abiotic stresses in horticulture (Gorbe *et al.*, 2004), hyperspectral imaging to predict the above-ground biomass of individual rice plants (Feng*et al*., 2013), 3D laser scanners to reconstruct and analyze deciduous saplings (Delagrange *et al*., 2011), PET to dissect dynamic changes in plant structure and function (Jahnke *et al*., 2009), MRI to analyze belowground damage to sugar beets (Hillnhutter *et al*., 2012), and X-ray imaging to quantify roots in soil (Flavel *et al*., 2012). However, little effort has been made in the dynamic phenotyping of rice tiller inner structures at high spatial resolution and high-throughput.

The rapid development of high-throughput phenotyping technology has accelerated the genetic mapping of important agronomic traits in crops. With the precision field phenotyping platform, QTLs (quantitative trait loci) for controlling biomass were identified in triticale (Busemeyer *et al*., 2013). The panicle-related image-analysis pipeline PANoram，promoted the genetic dissection of rice panicle traits (Crowell *et al*., 2014). With abundant genetic variations in natural populations, combinations of high-throughput phenotyping and genome-wide association studies (GWAS) have been conducted to reveal the natural genetic variation and to dissect the genetic architecture of complex traits, such as biomass, grain yield, leaf traits, panicle, and salinity tolerance (Yang *et al*., 2014; Yang *et al*., 2015; Al-Tamimi *et al*., 2016; Crowell *et al*., 2016).

Tiller numbers and angles are two key components of plant architecture that affect rice grain yield (Springer, 2010). Tiller number largely determines panicle number, a key factor in yield. Many tiller-related genes have been identified in recent years, such as *MOC1*(Li *et al*., 2003), *OsTB1*(Takeda *et al*., 2003), and *IPA1* (Jiao *et al*., 2010). These genes are involved in the initiation and outgrowth of axillary meristems and in the auxin and strigolactone signaling pathway that controls rice tillering (Li *et al*., 2003; Takeda *et al*., 2003; Guo et al., 2013). miRNAs are also involved in rice tillering by regulating the expression of target genes (Xia *et al*., 2012; Liang *et al*., 2014). *MOC1*, which was first isolated and characterized in the control of rice tillering, positively regulates tiller number by initiating axillary buds and promoting their outgrowth (Li *et al*., 2003). Tiller angle, which determines the plant density, has undergone domestication and improvement. Small tiller angles make plants more efficient in photosynthesis; therefore, dense planting is needed during rice cultivation (Yu *et al*., 2007). Several tiller-angle related genes, such as *TAC1, TAC3, OsLIC,* and *PROG1*, have been identified and characterized (Yu *et al*., 2007; Jin *et al*., 2008; Wang *et al*., 2008; Dong *et al*., 2016). *TAC1* is a major gene that positively controls tiller angle by forward genetics (Yu *et al*., 2007). A variant in the 3'-UTR changes the mRNA level, and higher mRNA levels contribute to a larger angle. Based on previous studies, nucleotide diversities in *TAC1* are low, and only one SNP in the coding region was found, resulting in synonymous substitution among 113 cultivated rice varieties. The small-angle allele of *TAC1* only exists in the *japonica* accessions (Jiang *et al*., 2012).

In the present work, we developed a high-throughput micro-CT-RGB (HCR) imaging system to extract tiller-related phenotypic traits with high spatial resolution (97 μm) and high efficiency (~310 pots per day). A rice panel containing 234 accessions were phenotyped non-destructively at 9 time points during the tillering process, and 730 traits were extracted by HCR and used to perform GWAS. Our results demonstrate that combining HCR and GWAS provides new insight into the genetic basis of rice tillering and plant architecture.

## Materials and Methods

### Plant material and experimental design

Considering the strong population differences between *indica* and *japonica* accessions and the high diversity in *indica* subpopulations (Huang *et al*., 2010), 234 *indica* accessions were used in our study. For each accession, one rice plant was detected by the HCR imaging system. The genotype information for the 234 accessions was retrieved from the website "RiceVarMap" (http://ricevarmap.ncpgr.cn/). The detailed information from the 234 rice accessions was obtained via the website (http://ricevarmap.ncpgr.cn/cultivars_information/). The seeds from the 234 rice accessions were sown in the field on 25 May 2015 and transplanted to pots on 16 June 2015. Each pot was filled with 5 kg soil (pH =5.45, total nitrogen: 0.241 g/kg, total potassium: 7.20 g/kg, total phosphorus: 0.74 g/kg, alkali-hydrolyzable nitrogen: 144.06 mg/kg, available potassium: 188.64 mg/kg, available phosphorus: 16.81 mg/kg, organic matter: 46.55 g/kg). During the tillering stage (41~67 days after sowing), the 234 rice accessions were automatically measured every three days and measured 9 times using HCR. After harvest, 203 rice plants were threshed and then inspected by YTS (yield traits scorer, Yang*et al.*, 2014) to extract grain yield. Thirty-five rice plants and standard plastic pipes were manually measured (Supplementary Fig. S1).

### Image acquisition of HCR

The control flow of image acquisition included the following steps (Supplementary Fig. S2): (1) the computer’s communication with the PLC and RGB camera was checked; (2) the X-ray flat panel detector was opened; (3) the working mode of the X-ray flat panel detector was selected; (4) the link with the X-ray flat panel detector was checked; (5) the mode information for the X-ray flat panel detector was retrieved; (6) the X-ray flat panel detector was used to grab images; (7) X-ray images and RGB images were obtained simultaneously; (8) X-ray images and RGB images were stored simultaneously; (9) X-ray image acquisition was stopped; (10) the X-ray flat panel detector link was closed; (11) the RGB camera and serial port were closed. The HCR image acquisition was implemented with LabVIEW 8.6 (National Instruments, US).

### Image analysis and traits extraction by HCR

Supplementary Fig. S3 and Supplementary Note S1-10 show the image analysis and trait extraction by the HCR system. Before image collection, the micro-CT system was off-set-calibrated and gain-calibrated. After calibration, the micro-CT system acquired 380 images while the rice plant rotated 360°. One row of X-ray projected images of the same height as the 380 X-ray projected images, was selected to form a sinogram, covering 380 orientations (step 0.6°, entire angle 0.6°×380, ~220°). Using the FBP algorithm and GPU acceleration technique, the inner structure of the rice tiller was reconstructed. By removing the small areas and regions with a predefined threshold, we counted 14 tiller traits, including tiller number, size and shape. Finally, when 2 transverse tiller images were reconstructed at 2 different heights (row 600 and row 650), 3 rice angle traits (mean, max, and standard deviation of the tiller angles) was calculated using the spatial location of the central point of the rice tiller images. Using the image analysis for the RGB images (Yang *et al*., 2014), 51 morphological features, 1 color trait, and 6 histogram features were calculated.

### Operation of the HCR

As shown in Supplementary Fig. S4 and Supplementary Fig. S5, the HCR operational procedure included the following steps: (1) the chiller was turned on and the water temperature maintained at 20℃; (2) offset calibration was performed; (3) gain calibration was performed; (4) one pot-grown rice plant was transported to the rotation platform; (5) the X-ray source was turned on and the inspection was started; (6) 380 CT images and 20 RGB images were obtained; (7) the next pot-grown rice plant was transported to the rotation platform; (8) when all the tasks were complete, the image acquisition software designed using LabVIEW was stopped.

### Growth modeling and yield predication using phenotypic traits

To test the prediction ability of the different models for TTA and TPA, 6 models, including linear, power, exponential, logarithmic, quadratic, and logistic, were built and compared. The modeling results were evaluated by comparing the R^2^, MAPE, and SDAPE values. The statistical analyses of the 6 TTA and TPA models (linear, power, exponential, logarithmic, quadratic, and logistic) were developed with LabVIEW 8.6 (National Instruments, Inc., USA). To evaluate the variance explained by the rice grain yield in the early growth stages, linear stepwise regression analysis was performed with the rice tiller traits using SPSS software (Statistical Product and Service Solutions, Version 13.0, SPSS Inc., USA).

### Genome-wide association study

A total of 2,863,169 single nucleotide polymorphisms (SNPs) with a minor allele frequency ≥0.05 were used for GWAS, and the number of accessions with minor alleles for the SNPs was more than 6. Information on these SNPs can be accessed from the ‘RiceVarMap’ database (http://ricevarmap.ncpgr.cn/). As in previous studies, the genome-wide significance threshold was set at 1.66×10^−6^ to control for false positives (Yang *et al*., 2015). A mixed-model approach with the factored spectrally transformed linear mixed models (FaST-LMM) program was used for the GWAS (Lippert *et al*., 2011). The kinship coefficient (K) values were defined as the proportion of identical genotypes for the 188,165 evenly distributed random SNPs (Xie *et al*., 2015). Lead SNPs for each trait were determined using the ‘clump’ function of Plink (Purcell *et al*., 2007). Potential candidate genes were obtained using the ‘clump-range’ function of Plink (Purcell *et al*., 2007). Considering the strong LD(linkage disequilibrium) of rice, a region in which the distance of adjacent pairs of associated SNPs was less than 300 kb was defined as the locus (Yang *et al*., 2015). Haplotypes were determined based on the significant genetic variants.

## Results

### High-throughput micro-CT-RGB phenotyping system (HCR)

The bi-modal imaging system, including micro-CT and RGB imaging, was developed to non-destructively extract 74 phenotypic traits synchronously. Among these 74 traits, tiller number, shape, area, and angle were extracted by CT images, and plant architecture, texture, and color traits, and digital biomass were extracted by RGB images. The definitions and abbreviations of the phenotypic traits are shown in Supplementary Table S1. The bi-modal imaging system consists of 9 main elements: an X-ray source (Nova600, OXFORD, UK), an X-ray source chiller (Nova600, OXFORD, UK), an X-ray flat panel detector (PaxScan 2520DX, VARIAN, USA), a RGB camera (AVT Stingray F-504B, Allied Vision Technologies Corporation, GER), a white light, a rotation platform (MSMD022G1U, Panasonic, Japan), a lead chamber, a computer (M6600N, Lenovo, CHN), and a PLC controller (CP1H, OMRON corporation, Japan) (shown in Fig. 1A, B). The configuration of the HCR system is provided in Supplementary Fig. S6, and shows that the CT system’s field of view (FOV) is 149 mm (height) × 186 mm (width) and the spatial resolution is 97 μm. The RGB imaging system’s FOV is 1607mm (height) × 1347 mm (width) and the spatial resolution is 656 μm. The main specifications of the HCR inspection unit are shown in Supplementary Table S2.

**Figure 1.**
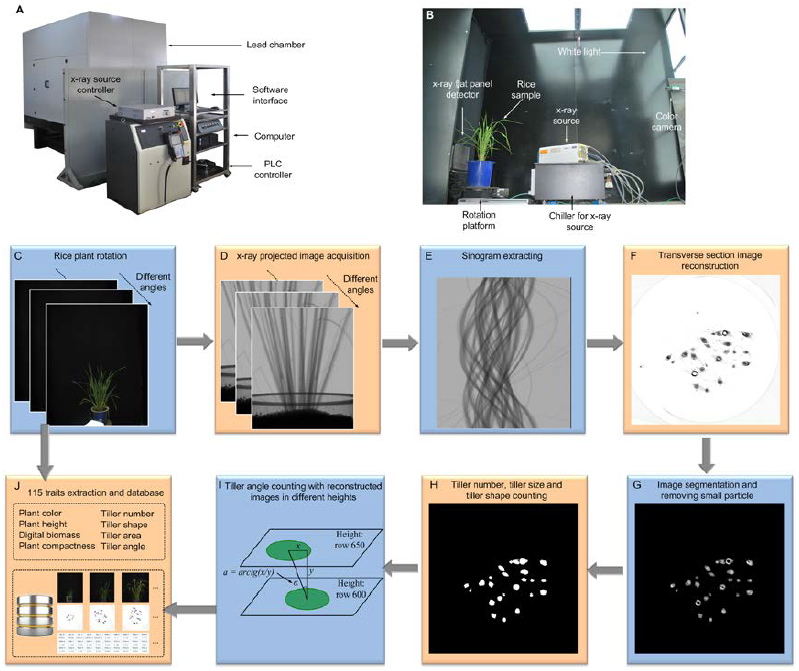
High-throughput micro-CT-RGB bi-modal imaging system. (A) The prototype of the micro-CT-RGB system and (B) layout of the inspection unit. The 78 rice shoot traits and 37 tiller traits were obtained via the following steps:(C) and (D) as the rice sample rotated, 20 color images and 380 X-ray projected images in different angles were acquired synchronously; (E) one row of X-ray projected images at the same height as the 380 X-ray projected images, which formed a sinogram, covering 380 orientations was selected (step 0.6°, entire angle 0.6°×380, ~220°); (F) conventional filtered back-projection (FBP) algorithm was applied to obtain the reconstructed transverse section image of rice tillers; (G) and (H) after image segmentation and removal of small particles, the tiller number, size and shape can be counted; (I) when 2 transverse tiller images were reconstructed at 2 different heights (row 600 and row 650), the rice angle was calculated using the spatial location of the central point of the rice tiller images; (J) 78 rice shoot traits (plant color, plant height,digital biomass, and plant compactness) and 37 tiller traits (tiller number, shape, area, and angle) were extracted and stored with the image analysis pipeline. A database was set up to collect RGB images, micro-CT images and phenotypic traits.

When the rice plant is rotated on the rotation platform (Fig. 1C), 20 color images and 380 X-ray projected images (Fig. 1D) in different angles are acquired synchronously. All phenotypic traits were obtained using the following steps: (1) one row of the X-ray projected image at the same height as the 380 X-ray projected images was selected to form a sonogram (Fig. 1E) covering 380 orientations (step 0.6°, entire angle 0.6°×380, ~220°); (2) a conventional filtered back-projection (FBP) algorithm was applied to obtain the reconstructed transverse section image of the rice tiller (Fig. 1F); (3) after image segmentation and small particle removal (Fig. 1G), tiller number, size and shape were counted (Fig. 1H); (4) when 2 transverse tiller images were reconstructed at 2 different heights (row 600 and row 650), the rice angle was calculated using the spatial location of the central point of the rice tiller images (Fig. 1I); (5) finally, 57 phenotypic traits, including plant color, plant height, digital biomass, and plant compactness, were obtained from the RGB images and analyses. A database, including the RGB and micro-CT images and the phenotypic traits, was set up (Fig. 1J). The reconstructed images of one rice sample (C055, Sanbaili) at different heights (10.7-54.3 mm distance from the soil surface) is shown in Supplementary Video S1. The image acquisition and analysis pipeline were developed using LabVIEW 8.6 (National Instruments, US), and the details were described in the Methods section.

As shown in Supplementary Fig. S4, the time taken for one CT image was 0.6 seconds, and 380 CT images were acquired for each plant; thus, approximately 228 seconds (0.6 seconds × 380) were required to complete the CT inspection of one pot-grown rice plant. The time taken for one RGB image was 0.6 seconds and 20 RGB images were acquired synchronously. The time taken for manual transfer is approximately 50 seconds. Therefore, when continuously operated for 24 hours each day, the HCR system’s total throughput is 310 pot-grown rice plants (~278 seconds per plant).

### Performance evaluation of tiller traits extraction

To evaluate the accuracy of the micro-CT unit, 8 plastic round pipes (fixed in one pot as shown in Supplementary Fig. S7) were measured manually by two people (phenotypic traits are shown in Supplementary Table S3) and automatically measured 10 times repeatedly by the micro-CT unit (phenotypic traits are shown in Supplementary Table S4). The mean absolute percentage error (MAPE) of the automatic versus manual measurements were 0.02~1.38%, 0~6.38%, and 0.12~1.87% for tiller diameter, stem wall thickness, and tiller angle, respectively (Fig. 2A). The computational formulas of MAPE were defined by Eqs. 1.

**Figure 2.**
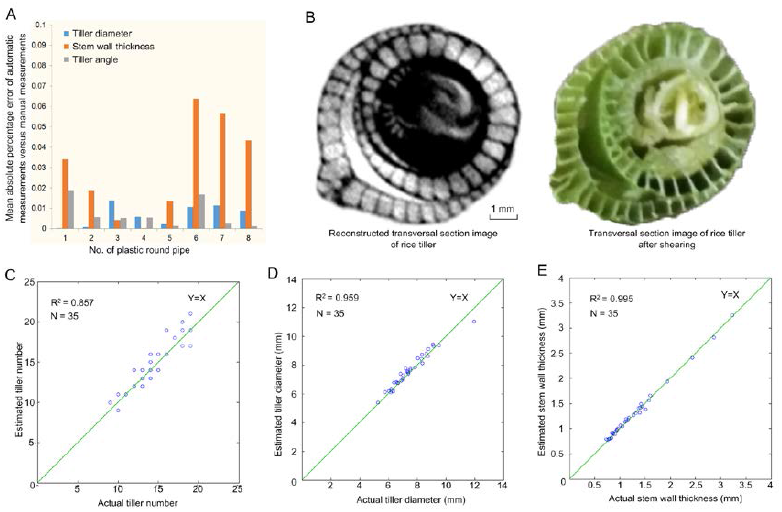
Comparison of results obtained via automatic measurements versus manual measurements. (A) The absolute percentage error of automatic measurements versus manual measurements of 8 round plastic pipes; (B) The reconstructed transverse section image of the rice tiller versus actual transverse section image of the rice tiller after shearing; Scatter plots of manual measurements versus automatic measurements with micro-CT unit for the tiller number (C), tiller diameter (D) and stem wall thickness (E).

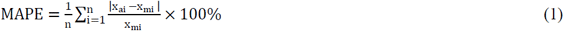

To evaluate the reconstruction quality of the rice tiller, a reconstructed transverse section image (spatial resolution of 30 μm) using micro-CT and its actual transverse section image after shearing are shown in Fig. 2B. In addition, there was a trade-off between the CT image resolution and CT scan area. To scan all the rice tillers, the spatial resolution was set at 97 μm and the FOV of the CT system was 149 mm (height) × 186 mm (width) (Supplementary Fig. S6). Next, 35 rice plants (Supplementary Table S5) were measured both automatically and manually (repeat twice) to verify the measuring accuracy using micro-CT. The R^2^ values of the manual measurements versus automatic measurements were 0.857, 0.959, and 0.995 for tiller number, tiller diameter, and stem wall thickness, respectively (Fig. 2C-E).

### Phenotyping database extracted by HCR at 9 time points

During the tillering stage, 234 rice plants were automatically measured by HCR at 9 different development time points (once every 3 d, starting from 41 ~ 67 d after sowing). All the phenotypic data and images can be viewed and downloaded via the link http://plantphenomics.hzau.edu.cn/checkiflogin_en.action and then following these steps: (1) select ‘rice’; (2) select ‘2015-tiller’ in the year section; (3) select one of the accession IDs in the ID section and then press ‘search images’; (4) 9 CT images and 9 side-view color images can be viewed and downloaded; (5) a similar process can be used to view and download phenotypic traits by pressing ‘search data’. The detailed procedure for the database is shown in Supplementary Fig. S8.

### Screening the dynamic process of rice growth at the tillering and jointing stages

After all phenotypic images and data were obtained for the 9 time points, we screened the dynamic process of the rice growth and determined the most active tillering and initial jointing stages. As illustrated in Fig. 3A-I, 9 side-view RGB images and 9 reconstructed images for each rice plant were obtained for the following image analysis. The red circle in Fig. 3B-E shows the dynamic tillering and jointing processes. At the second time point (Fig. 3B), the first pith cavity appeared, indicating that this plant progressed into the jointing stage. As illustrated in Fig. 3J, from the dynamic change of the first derivative of the total tiller area, we can determine the most active tillering stage, as indicated by the blue arrow with the maximum value of the first derivative of the total tiller area. The tiller growth of the rice plant during the first 6 periods was relatively faster than that of the later periods. Similarly, from the number change of the rice accessions in the initial jointing stage, we can see that the initial jointing stage was accompanied by the most active tillering stage (Fig. 3K). Interestingly, the growth curve of the GCV (green color value) before the 5^th^ time point indicates that the GCV value became smaller (indicating more dark green leaves with greater nitrogen content), and after the 5^th^ time point, the GCV value became larger (indicating more light green leaves with less nitrogen) (Fig. 3L). As illustrated in Fig. 3M, from the dynamic change of first derivative of the mean tiller angle, we see that the tiller angle showed little change during the tillering stage.

**Figure 3.**
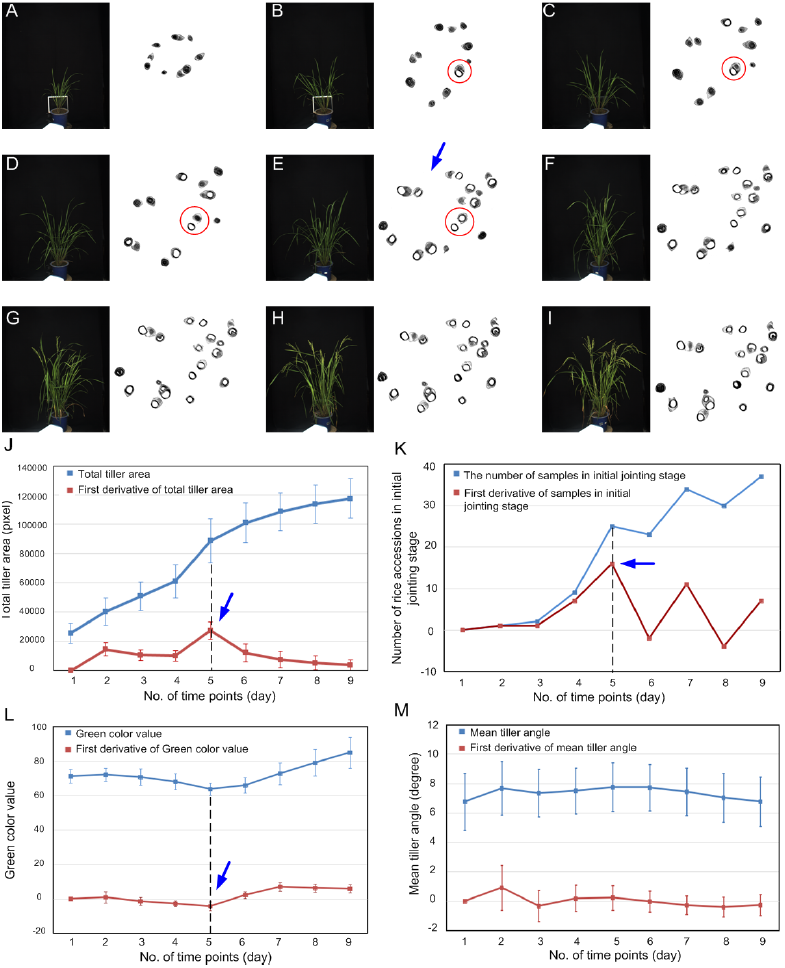
Screening the dynamic process of rice growth at the tillering stage and jointing stage. (A-I) The RGB images and reconstructed CT images at 9 different growth time points; (J) diagram of total tiller area and first derivative of total tiller area; (K) diagram of sample numbers in initial jointing stage and first derivative of sample numbers in jointing stage; (L) diagram of green color value and first derivative of green color value; (M) diagram of mean tiller angle and first derivative of mean tiller angle. The error bars represent the standard deviation between the accessions.

In addition, the dynamic growth curves of 27 representative traits for the tiller and the entire plant are presented in Supplementary Fig. S9. The first derivation of H (plant height), W (plant width), and TPA (total projected area of the rice plant) reached the highest value at the 5^th^ time point, supporting the previous result that the plant growth reached the highest speed in the active tillering stage (5^th^ time point). The dynamic growth of one rice accession (C055, Sanbaili) is shown in Supplementary Video S2.

### Predication of tiller growth and digital biomass accumulation

It would be helpful if we could design a growth model using the phenotypic data obtained in the early growth stage to predict the final digital biomass. In our previous study, total projected area (TPA) was correlated with actual biomass (Yang et al., 2014). Beyond the manual tiller number count, the total tiller area (TTA) extracted by micro-CT can quantify tiller growth more accurately than the tiller number. Fig. 4A, B show the heatmaps of TTA and TPA for the 234 accessions at 9 different time points. Here, we tested 6 models (linear, power, exponential, logarithm, quadratic, and logistic models) of TTA and TPA at the 9 points. The results were evaluated by comparing R^2^, MAPE, and the standard deviation of the absolute percentage error values (SDAPE). As shown in Supplementary Table S6, the logistic models of TTA and TPA showed slightly better prediction ability (the R^2^ was 0.969 and 0.985, the MAPE and SDAPE were both below 6.5%). The actual results versus predicted results of the TTA and TPA are shown in Fig.4C and Fig.4D, respectively.

**Figure 4.**
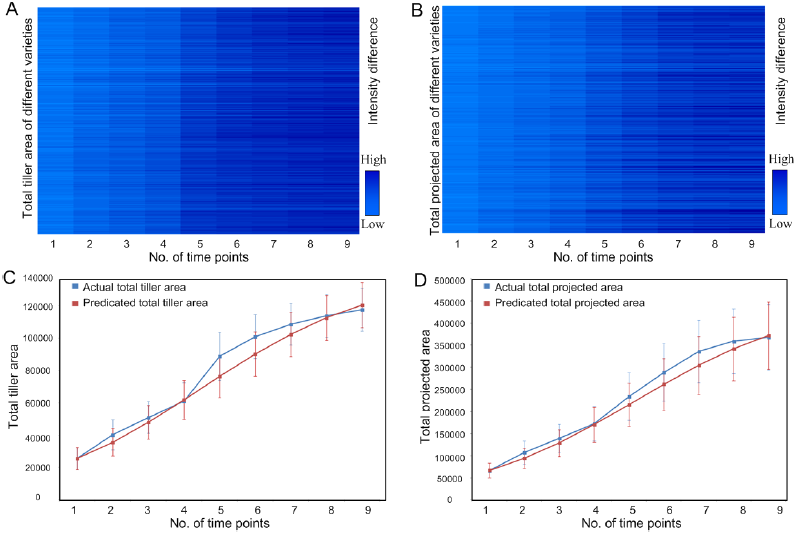
Heatmap and prediction of total tiller area growth and total protected area growth. (A-B) Heatmap of total tiller area (TTA) and total protected area (TPA)of the 234 individuals at 9 different time points; (C)comparison of actual total tiller area (blue line) and predicted total tiller area (red line); (D) comparison of actual total projected area (blue line) and predicted total projected area (red line).Error bars represent the standard error of the TTAorTPA of 234 samples at each time point.

### Predication for rice grain yield and shoot dry weight in the early growth stage

It would benefit rice breeding if we could use the automatically measured phenotypic traits, particularly the traits measured in the early development stages, to predict the final grain yield and shoot dry weight. The R value distribution for modeling grain yield in the 9 different tillering stages is shown in Fig. 5A, which shows that by adding the total tiller area (TTA), the R range increased from 0.30-0.41 to 0.35-0.51, particularly at the 5^th^ time point. After the 5^th^ time point, nonfertile tillers began to grow, providing a possible explanation why the R value decreased. Fig. 5B showed that the modeling accuracy for the shoot dry weight is improved by adding total tiller area. Moreover, we also compared the correlation between TN, TTA and grain yield. The R value of TN_5 versus grain yield was 0.094 (Fig. 5C), and the R value of TTA_5 versus grain yield was 0.512 (Fig. 5D).

**Figure 5.**
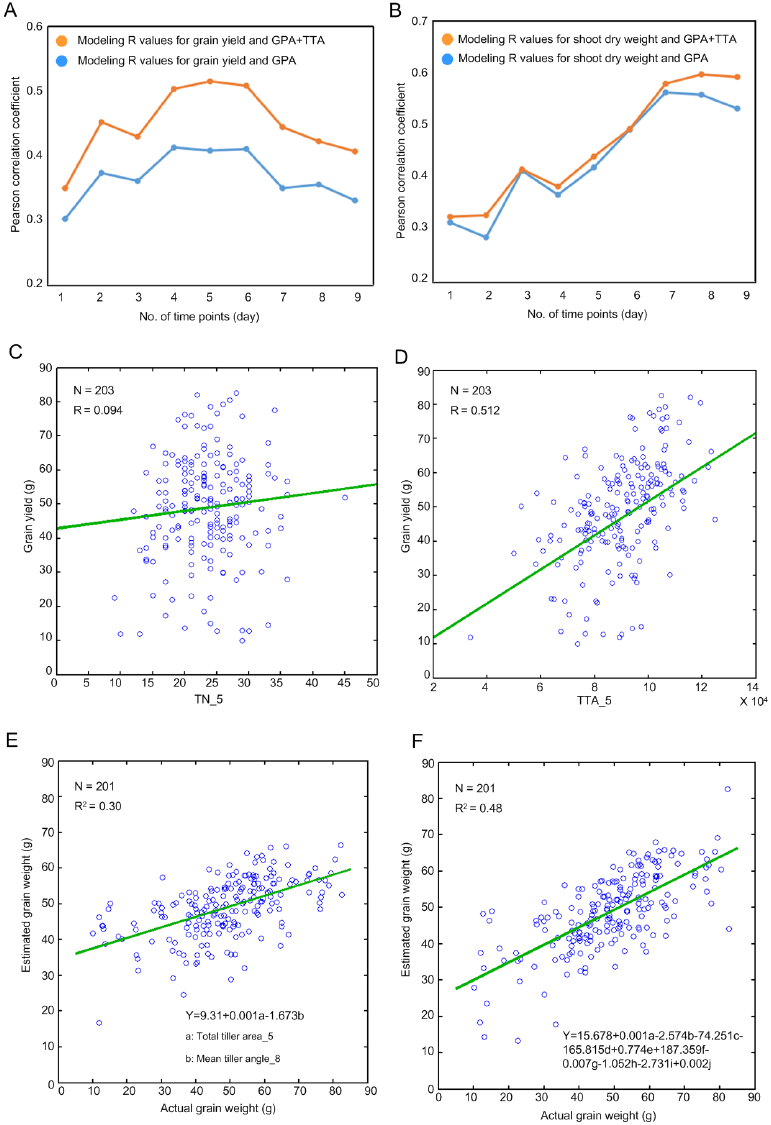
Predication of grain yield and shoot dry weight. (A) the modeling accuracy change for grain yield at 9 time points; (B) the modeling accuracy change for shoot dry weight at 9 time points; (C) the scatter plot of tiller number versus grain yield at the 5^th^ time point; (D) the scatter plot of total tiller area versus grain yield atthe 5^th^ time point; the scatter plot showing the relationship between the actual grain yield and estimated grain yield using the predicted formula by (E) 2 traits and (F) 10 traits; a, b, c, d, e, f, g, h, i and j represent TTA_5, MEANTA_8, THR_4, FDIC_7, MAXTAPR_7, FDIC_8, SDTTA_5, TN_3, MEANTAPR_2 and MAXTTA_2, respectively.

When only 2 phenotypic traits were selected, 30% of the grain yield variance was explained (Fig. 5E). The two phenotypic traits were both tiller traits, which included TTA_5 (total tiller area measured at the 5^th^ time point) and MEANTA_8 (mean value of the tiller angle measured at 8^th^ time point). We found that the rice yield can be increased by higher TTA_5 and lower MEANTA_8. Up to 48% of the grain yield variance can be explained by combining 10 traits across all 9 time points (Fig. 5F). As shown in Supplementary Fig. S10, the R^2^ value range from 0.34 to 0.46 by combining from 3 traits to 9 traits.

### Genome-wide association study

We performed GWAS of 732 traits (including 730 traits measured by micro-CT-RGB, yield and biomass) and identified 402 significantly associated loci (Supplementary Data S1). In total, 182 and 332 loci were associated with traits measured by micro-CT and RGB, of which 70 and 220 were exclusively detected by micro-CT and RGB respectively. The numbers of loci associated with traits of different time points were different, ranging from 61 and 87. For example, the numbers of loci of time point 1, 5, 9 were 61, 86, 69; the numbers of overlapped loci of T1 and T5, T5 and T9, T1 and T9 were 14, 17, 8; only 4 loci were detected at all the three time points (Fig. 6A). Of 402 loci, 353 and 135 loci were detected by the micro-CT-RGB traits of nine time points and the derived growth-rate related traits, and 86 loci were simultaneously detected by the two kinds of traits. Of the 353 loci, 191 loci were only detected at one time point while other loci were detected at not less than two time points; only one locus on chromosome 9 (locus 302) were detected at nine time points (Fig. 6B). Further we found the locus were significantly associated with MEANTA (mean of multiple-tiller angles for a plant) measured by micro-CT (Fig. 6C), suggesting the locus could control tiller angle. These results demonstrate the existence of dynamic and static genetic components during rice growth stage.

**Figure 6.**
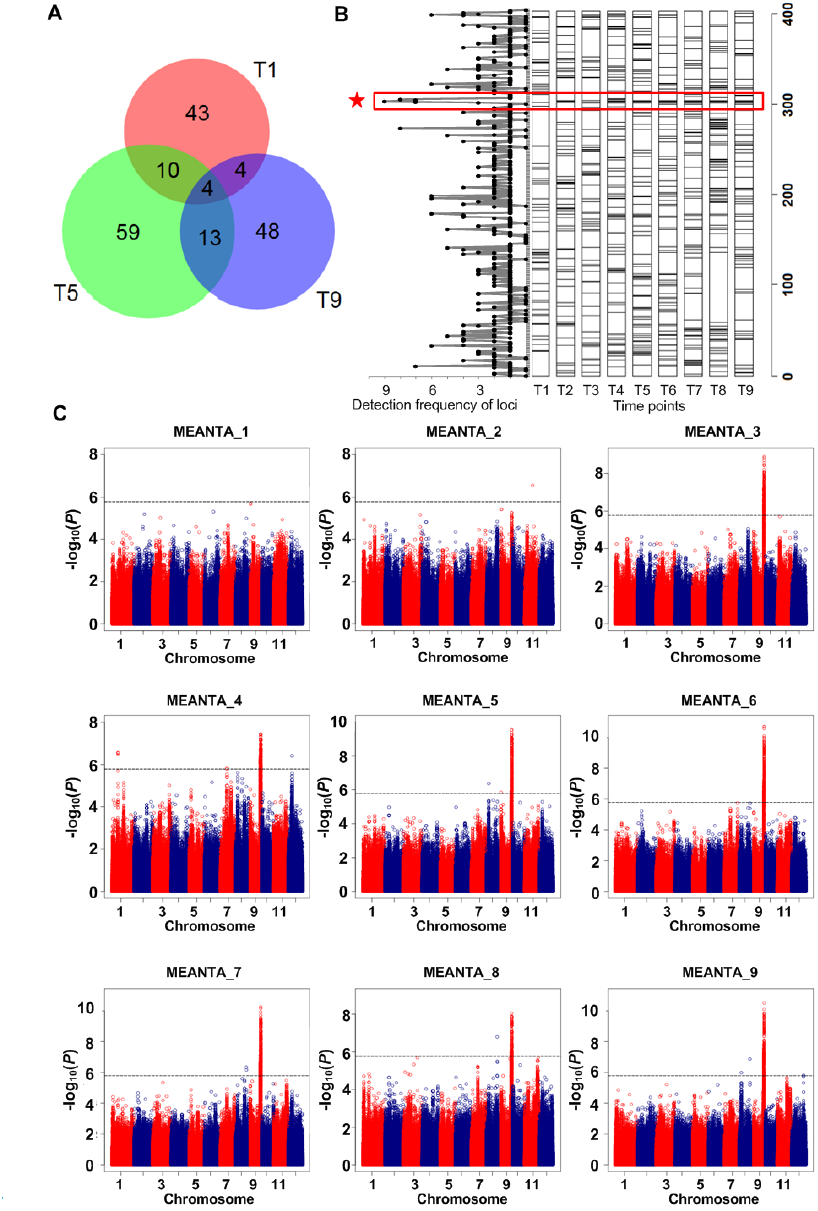
GWAS results of traits of nine time points measured by micro-CT-RGB. (A) Venn diagram showing number of associated loci at the time point 1, 5, and 9. (B) The frequency and distribution of loci associated with traits at nine time points (T1-T9). (C) GWAS plots of MEANTA (mean of tiller angles) of nine time points. The strongest association signal on chromosome 9 corresponded to the locus of highest detection frequency.

For the locus 302, LD decayed slowly (r^2^=0.57 between SNPs sf0920227209 and sf0920733864) in a 500 kb-region. *TAC1*, the cloned gene controlling tiller angle (Yu *et al*., 2007), was located at the locus. We found 15 significant SNPs distributed in the 3'-UTR region, coding region, and 1 kb promoter region and a significant 1-bp indel in the 3'-UTR region (Fig. 7A). All the SNPs in the coding region caused synonymous mutations. Consistent to a previous study (Yu *et al*., 2007), the variants in the 3'-UTR caused the mRNA level polymorphisms, resulting in the tiller angle diversity. Three haplotypes for the gene were found in our association mapping panel. Tiller angles were significantly different among them (*P*=5.15E-07, ANOVA) and those of rice accessions containing haplotype H3 were much smaller (Fig. 7B). Minghui 63 (a known restorer line) and Zhenshan 97 (a known maintainer line) contained haplotype H2 and H3, respectively (Fig. 7C).

**Figure 7.**
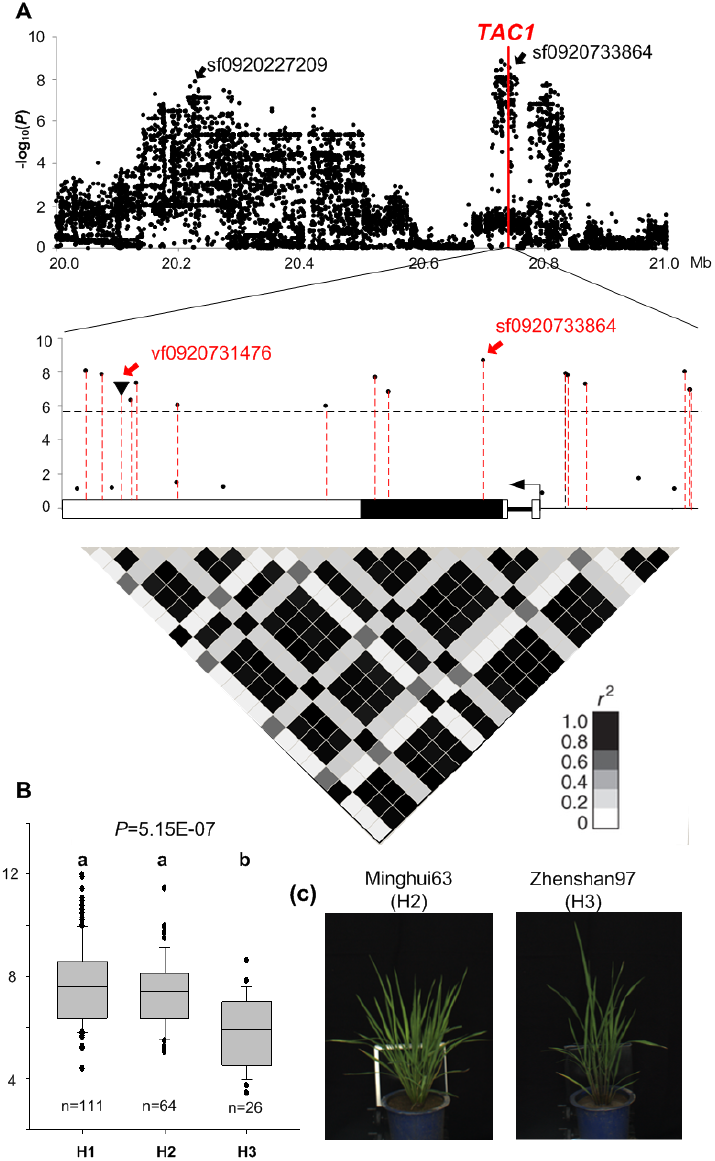
Association analyses of *TAC1*and MEANTA_3. (A) Local Manhattan plots and heat map showing LD level of *TAC1* region. (B) Haplotype analyses of *TAC1. P* value was calculated by ANOVA. Multiple-haplotype comparison was conducted using LSD method and different letters above box plot indicated significant difference. Images of two representative varieties-Minghui63 (from H2 haplotype group) and Zhenshan97 (from H3 haplotype group).

Further, we found two loci containing associations with both micro-CT-RGB traits and yield. A lead SNP sf0401216812 on chromosome 4 was associated with AGRTTA_5 indicating growth rate of tillering at the 5^th^ time point (*P*MLM=1.16E-05) and yield (*P*MLM=8.40E-04), and genotype G at the SNP site corresponded to the superior allele for the two traits (Fig. 8A). Another lead SNP sf0630983585 on chromosome 6 was associated with AGATPA_4 indicating growth rate of shoot weight at the 4^th^ time point (*P*MLM=1.14E-06) and yield (*P*MLM=2.93E-04), and genotype G at the SNP site corresponded to the superior allele for the two traits (Fig. 8B). The favorable alleles of the two loci were minor alleles and would be beneficial for rice high-yield breeding. These results indicate that the vigor of rice plant during tillering stage contributes to the final yield.

**Figure 8.**
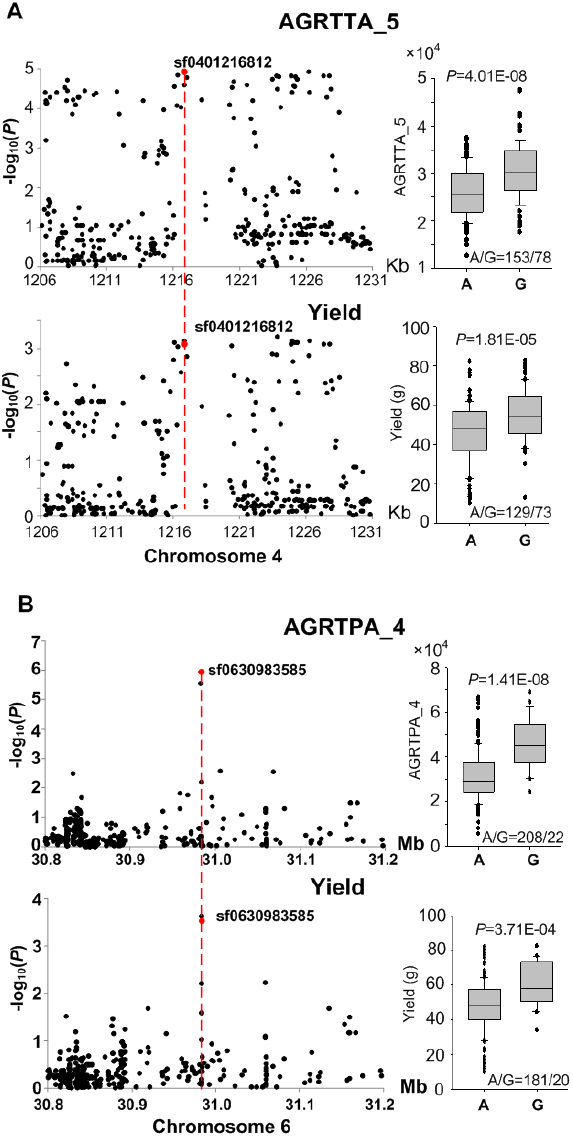
Co-localized loci associated with traits measured by micro-CT-RGB and yield. (A) The locus on chromosome 4 associated with AGRTTA_5 measured bymicro-CT and yield. (B) The locus on chromosome 6 associated with AGRTPA_4 measured by RGB and yield.

## Discussion

The traditional methods of determining rice tiller traits are destructive, labor-intensive, and time-consuming. Micro-CT, a computed tomography technique originally developed for structural imaging of small animals (Yang *et al.*, 2010), can also be an option for examining the inner structure of rice plants with multiple tillers. In addition, by developing an image analysis pipeline, the HCR system can non-destructively extract rice phenotypic traits and provide plant growth data in vertical and horizontal dimensions. Compared to traditional rice tiller phenotyping, HCR has the following advantages. (1) The 3D spatial location can be obtained by CT, thus, some traits, such as tiller angle, can be extracted with more accuracy rather than manually measuring them with a protractor, as shown in Fig. 1I. (2) The CT system can be easily integrated with an RGB imaging device, allowing more traits (total of 74 traits) to be extracted simultaneously. (3) The time needed for acquiring the projected CT image of one plant is approximately 278 seconds, and the time required for extracting subsequent traits is approximately 120 seconds combined with GPU acceleration, thus improving the measuring efficiency per plant. (4) Many novel traits, such as TTA and TPA, can be investigated with the bi-modal imaging system at different time points. In comparing the tiller number and grain yield, the total tiller area (TTA) had a better correlation with grain yield and provided a better quantification of tiller growth (Fig. 5C and 5D). Finally, (5) these new dynamic traits in plant growth and tiller development can dissect the genetic mechanisms involved in rice growth.

With numerous traits extracted by HCR, GWAS detected many significant association signals. The number of loci detected at different time point was different. Some loci were identified at a specific time point while other loci were identified at multiple time points, indicating the dynamic and static genetic components during rice growth stage. Only one locus on chromosome 9 related to tiller angle was scanned at 9 time points and a priori gene *TAC1* was located at the locus. Six significant SNPs and a significant INDEL were enriched in 3'-UTR region. We observed three major haplotypes for the gene in our association mapping panel and significant difference of tiller angle among the three haplotypes. Although most *indica* accessions harbored the haplotype of the wider tiller angle for *TAC1,* some *indica* accessions harbored the haplotype of the narrow tiller angle, which was not found in previous studies. The polymorphisms in *TAC1* can be further developed into markers for breeding selection for density planting. Co-localized loci between HCR traits indicating vigor of rice plant during growth stage and yield were found, and HCR traits had higher detection power than yield. The superior alleles of the loci were minor alleles, which would be used for breeding of high yield.

## Conclusions

In this study, we developed a high-throughput micro-CT-RGB (HCR) imaging system to extract tiller-related phenotypic traits with high spatial resolution (97 μm) and high efficiency (~310 pots per day). A rice panel containing 234 accessions was phenotyped non-destructively at 9 time points during the tillering stage, and totally 730 traits were extracted by HCR and used to perform a GWAS. A total of 402 significantly associated loci were identified by GWAS, and dynamic and static genetic components were found across the nine time points. A major locus associated with tiller angle was detected at nine time points and a priori gene *TAC1* was located at the locus. Significant variants associated with tiller angle (evaluated by MEANTA) were enriched in the 3'-UTR of *TAC1.* Three haplotypes for the gene were found and tiller angles of rice accessions containing haplotype H3 were much smaller. Further, two loci contained associations with both HCR traits and yield and the superior alleles were minor alleles, which would be beneficial for breeding of high yield and dense planting.

## Supplementary Data

Supplementary data are available at *JXB* online.

**Fig. S1.** Experimental design.

**Fig. S2.** Control flow of image acquisition.

**Fig. S3.** Diagram of image processing and feature extraction.

**Fig. S4.** Sequence diagram of micro-CT-RGB phenotyping system.

**Fig. S5.** Workflow chart.

**Fig. S6.** The configuration of micro-CT-RGB system.

**Fig. S7.** Plastic round pipes.

**Fig. S8.** Workflow chart of database.

**Fig. S9.** Dynamic growth curve of rice.

**Fig. S10.** Modeling results of grain yield.

**Note S1.** The source code of sinogram.

**Note S2.** The source code of computed tomography reconstruction.

**Note S3.** The source code of particle extraction.

**Note S4.** The source code of particle rotation.

**Note S5.** The source code of tiller diameter.

**Note S6.** The source code of tiller angle.

**Note S7.** The source code of fill holes.

**Note S8.** The source code of area traits.

**Note S9.** Color component extraction.

**Note S10.** Definition of the features.

**Table S1.** Abbreviation of 17 tiller traits, 32 tiller growth traits, 1 plant color trait, 2 digital biomass, 33 plant architecture traits, 21 texture traits, 16 digital biomass accumulation traits, 16 height accumulation traits, and 2 yield traits.

**Table S2.** Main specifications of micro-CT-RGB inspection unit.

**Table S3.** Manual measurements of 8 plastic pipes with 2 workers.

**Table S4.** Automatic measurements of 8 plastic pipes with 10 replications.

**Table S5.** Comparison of rice with automatically measured and manually measured.

**Table S6.** The Comparison of actual TTA/TPA and predicated TTA/TPA with 6 models.

**Video S1.** The reconstructed images of one rice sample at different heights.

**Video S2.** The dynamic growth of one rice accession.

**Data S1.** GWAS results.

## Acknowledgements

This work was supported by grants from the National Key Research and Development Program of China (2016YFD0100101-18), the National Natural Science Foundation of China (31770397), and the Fundamental Research Funds for the Central Universities (2662017PY058).

